# Identifying the spatial and temporal dynamics of molecularly-distinct glioblastoma sub-populations

**DOI:** 10.1101/2020.04.08.024091

**Authors:** Bethan Morris, Lee Curtin, Andrea Hawkins-Daarud, Matthew E. Hubbard, Ruman Rahman, Stuart J. Smith, Dorothee Auer, Nhan L. Tran, Leland S. Hu, Jennifer M. Eschbacher, Kris A. Smith, Ashley Stokes, Kristin R. Swanson, Markus R. Owen

## Abstract

Glioblastomas (GBMs) are the most aggressive primary brain tumours and have no known cure. Each individual tumour comprises multiple sub-populations of genetically-distinct cells that may respond differently to targeted therapies and may contribute to disappointing clinical trial results. Image-localized biopsy techniques allow multiple biopsies to be taken during surgery and provide information that identifies regions where particular sub-populations occur within an individual GBM, thus providing insight into their regional genetic variability. These sub-populations may also interact with one another through a competitive or cooperative nature; it is important to ascertain the nature of these interactions, as they may have implications for responses to targeted therapies. We combine genetic information from biopsies with a mechanistic model of interacting GBM sub-populations to characterise the nature of interactions between two commonly occurring GBM sub-populations, those with EGFR and PDGFRA genes amplified. We study population levels found across image-localized biopsy data from a cohort of 25 patients and compare this to model outputs under competitive, cooperative and neutral interaction assumptions. We explore other factors affecting the observed simulated sub-populations, such as selection advantages and phylogenetic ordering of mutations, which may also contribute to the levels of EGFR and PDGFRA amplified populations observed in biopsy data.

## 1 Introduction

Glioblastomas (GBMs) are the most common type of primary brain tumour occurring in adults and are a particularly aggressive form of cancer. Standard treatment consists of surgically removing as much of the tumour as thought safe, followed by radiation and the chemotherapy temozolomide [1]. In spite of such aggressive treatment regimens, GBMs inevitably recur, with median survival times remaining at just 14.6 months [2].

A major factor that is thought to contribute to treatment failure is the heterogeneous nature of GBMs. These tumours are known to be comprised of multiple sub-populations of cancerous cells, each with different genetic and phenotypic features [3]. Further to this, the particular sub-populations present in a tumour vary both between patients as well as within individual tumours, creating a disease characterised by both inter- and intra-tumoural heterogeneity. In addition to this genetic heterogeneity, a similar heterogeneity in response to treatment is observed among patients, with some patients responding to a particular treatment well and others not so. Furthermore, treatment response variability is also observed even within some individual tumours, with responses varying from one region of the tumour to another. These heterogeneities in treatment responses are thought to be at least partly due to the underlying sub-populations present within each individual tumour and their distribution throughout the tumour region, with individual sub-populations responding to a given therapy differently. Furthermore, the presence of such genetically and phenotypically distinct sub-populations may be the cause of failure of some therapies, particularly targeted therapies, since resistant clones may pre-exist within the tumour or non-resistant cells could survive through cooperation with other tumour cells [4, 3, 5, 6]. For this reason, gaining further understanding of this genetic heterogeneity and the implications for therapies will help to identify subgroups of patients with better responses to treatments and new targets for novel therapies.

Over recent years, advances in biopsy sampling techniques have been helping to characterize this inter- and intra-tumoural genetic heterogeneity exhibited by GBMs. Treatment decisions are based on the biomarkers present within a single biopsy specimen, which may only be representative of a small part of the tumour. This means that clinical decisions made based on this knowledge may not be the optimal therapeutic strategy to target the majority of cancerous cells composing the rest of the tumour, which may have different genetic features and respond better to another treatment option. Therefore, in order to gain more understanding of the genetic heterogeneity across the tumour region, image-localized multiple biopsy sampling techniques have been developed, allowing surgeons to collect multiple tissue samples from an individual during surgery and record information about the location in the tumour from where the sample was taken. Subsequent tissue analysis then identifies how the genetic profile of these samples differs from one region of the tumour to another, providing spatial and genetic information that gives insight into the inter- and intra-tumoural heterogeneity present in these complex tumours; examples of such techniques being employed can be found in [7], [8] and [9].

While the exact mechanism by which such genetic heterogeneity arises in GBMs is currently unknown, several possible theories have been proposed to explain this. These include the theory of clonal evolution, where tumours are thought to evolve through a process of acquiring mutations and natural selection; the cancer stem cell (CSC) model, in which a small population of CSCs give rise to and maintain the tumour through self-renewal and producing phenotypically diverse daughter cells; or possibly some complementary combination of these two theories [3]. In addition to understanding how genetic heterogeneity arises in GBMs and other types of cancer, further understanding of how such heterogeneity is maintained is also needed. One suggested mechanism of heterogeneity maintenance is called interclonal cooperativity, where interactions between different tumour cell populations are thought to be important; the theory suggests that some cells may acquire mutations that result in the promotion of other tumour cell sub-populations in some way [3, 10]. One consequence of this could be that a small population of genetically-distinct tumour cells plays an important role in tumour progression and, thus, targeting such a sub-population of cells may have additional negative effects on the rest of the tumour cell population. Therefore, identifying such genetically distinct tumour cell sub-populations and understanding their interplay with other cell populations may have important implications for the success of therapies.

Two such populations of interest are those with amplification of the Epidermal Growth Factor Receptor (EGFR) and the Platelet-Derived Growth Factor Receptor Alpha (PDGFRA) genes, i.e. cells with an increased number of copies of the genes encoding each protein. While amplification status relates to the number of copies of a particular gene that a cell has in its DNA, its *copy number aberration*, it also induces overexpression of these genes in tumour cells [11]. The EGFR and PDGFRA proteins are both members of the Receptor Tyrosine Kinase (RTK) family of cell surface receptors which bind to a variety of growth factors, cytokines and hormones and play a crucial role in the regulation of the signalling that controls cell proliferation, metabolism and survival [12]. Specifically, EGFR is a receptor that, upon binding, results in the activation of pathways that lead to cell proliferation, DNA synthesis and the expression of certain oncogenes [13] and its amplification has been shown to promote invasion in GBMs [14, 15] and be an unfavourable predictor for patient survival [16]. Meanwhile, PDGFRA is a receptor that, when bound, activates signalling pathways that promote oncogenesis [17, 18]. Due to the prevalence of EGFR and PDGFRA amplified tumour cells in GBMs, occurring in 43% and 11% of GBM samples in The Cancer Genome Atlas (TCGA) database, respectively [19], these sub-populations have become prime molecular targets for therapies and a number of inhibitor drugs have been developed for this purpose [5]. These therapies targeting EGFR and PDGFRA amplified cells, however, have had limited success in GBMs in clinical trials so far [5].

Several possible mechanisms of chemoresistance to these drugs in GBMs are discussed by Nakada et al. [5]. However, one possible mechanism of chemoresistance to EGFR and PDGFRA targeted therapies of interest is through the interaction of cell sub-populations with amplification of these genes; these cells may interact in a cooperative way that facilitates their survival or, conversely, competitively, such that the targeting of one population with therapy benefits the other by removing its competitor. While the interactions between EGFR and PDGFRA amplified sub-populations are currently not well understood, it has been suggested that these cell populations may be interacting in a cooperative manner. For example, in experiments by Szerlip et al. [6] a form of cooperativity was observed between these cell populations, as combined inhibition of both receptors was needed to block activity of the PI3 kinase pathway- a pathway involved in the regulation of cell proliferation, apoptosis and migration [20]- in a mixed population of EGFR and PDGFRA amplified cells *in vitro*. In addition to this, Snuderl et al. [21] observed coexistence of these amplified sub-populations and suggested that they may co-evolve with similar fitness levels rather than compete during tumour evolution; the authors further suggest the possibility that these sub-populations cooperate to achieve a higher fitness level than each of the sub-populations individually [22, 21].

As stated previously, if EGFR and PDGFRA amplified sub-populations are indeed cooperating in GBMs, this will have implications for therapies targeting these cells and so it is important to gain more understanding of the nature of any interactions. In this paper, we present information from image-localized biopsies that provides insight into the distribution and co-occurrence of EGFR and PDGFRA amplified sub-populations throughout GBM tumours in a cohort of patients. While this provides important genetic and spatial information, this information is static and so it is difficult to extract any dynamic information that may help to identify the type of interactions occurring between these populations of tumour cells. Mathematical modelling could be a useful tool in this scenario, as models can be used to enhance current knowledge and provide deeper insight into complex biological processes. Therefore, we propose a novel mathematical model of interacting GBM sub-populations, where we investigate the effects of different interaction assumptions, namely cooperative, competitive and neutral (no) interactions, on the population level occurrence of EGFR and PDGFRA amplified cells *in silico*. We study population levels found across the image-localized biopsy data from a cohort of patients and compare this to model outputs under these different interaction assumptions. We explore additional factors affecting the patterns observed in our computational simulations, such as selection advantages and phylogenetic ordering of mutations, which may also contribute to the levels of EGFR and PDGFRA amplified populations observed in biopsy data. Finally, we discuss our results and the insight they provide into the evolution of these biologically complex tumours as well as our planned future work.

## 2 Materials and methods

### 2.1 Image-localized biopsies and tissue analysis

Patients with clinically suspected GBM undergoing preoperative MRI for surgical resection were recruited and the absence of previous treatment was confirmed. Institutional review board approval was obtained, along with written and informed consent from each participant prior to enrollment. During surgery, each neurosurgeon collected an average of 4–5 tissue specimens from each tumour and typically selected targets separated by ≥1 cm from different regions of the tumour based on clinical feasibility (e.g. accessibility of the target site, overlying vessels, areas of the brain that directly control function). The location of each biopsy was also recorded by neurosurgeons to allow for subsequent co-registration with multiparametric magnetic resonance imaging (MRI) datasets. More detail of the biopsy collection protocol can be found in [7].

To determine whether a biopsy sample contains tumour cells with the EGFR and PDGFRA genes amplified, copy number aberration (CNA) values associated with these genes were determined for all tissue samples using array comparative genomic hybridization (aCGH) as detailed in references [7, 23, 24]. Each tissue sample was then classified as being amplified in a given gene if the corresponding CNA value was greater than a given threshold and not amplified in that gene when below or equal to that threshold. Each biopsy sample, however, is likely to contain a mixture of healthy non-cancerous cells and tumour cells without and with varying degrees of gene amplification. Thus, the CNA value will be based on a mixed signal from a sample containing a mixture of cells with potentially different numbers of copies of the genes of interest and it is unclear what an appropriate threshold should be to determine the gene amplification status, which is a topic widely discussed in the literature [11, 25].

In this work, we choose to use a CNA threshold of 2.2; this threshold is chosen based on some prior knowledge and some assumptions about the levels of EGFR and PDGFRA amplification that we expect to see in our tissue samples. Firstly, diploid cells that are not EGFR or PDGFRA gene amplified will have an associated CNA value equal to 2 [11, 25], which applies to the healthy cells and non-amplified tumour cells in the tissue samples. Secondly, we assume that the EGFR and PDGFRA amplified cell sub-populations are homogeneous with respect to their gene copy numbers and, therefore, all cells in each of these sub-populations have the same CNA value associated to each of these genes, which we choose to equal 4; this corresponds to each of the alleles in an EGFR amplified cell containing an extra copy of the EGFR gene and similarly for the PDGFRA amplified population. Finally, since the neurosurgeons collect biopsy samples from various regions of each tumour, including the invasive edge where tumour cell density is low, we choose the CNA threshold in a way that will be sensitive to such low densities of EGFR and PDGFRA amplified cell sub-populations; we choose this low density threshold to be 10% of the tissue in a sample and assume that if a biopsy sample consists of 10% or less of either amplified cell sub-population, then the signal will be too low to be detectable in the corresponding CNA value and this sample will not be classed as being amplified in this gene. Therefore, the CNA threshold of 2.2 is derived as follows:

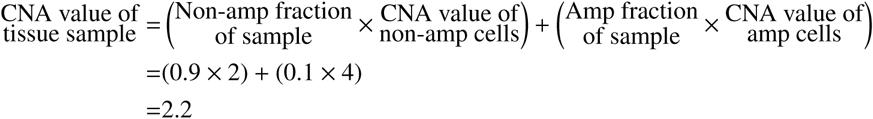

We note that tumour cells can exhibit varying degrees of gene amplification; for example, an EGFR amplified cell sub-population is likely to consist of cells containing a variety of copy numbers of the EGFR gene, with cells containing more than 100 copies in some cases [11]. This means that using a low CNA threshold to determine gene amplification status of a tissue sample in this way, could classify a sample containing a very low fraction of “highly amplified” cells as being gene amplified. However, we expect such cases to be rare and choose this CNA threshold to avoid excluding samples that contain cells that are amplified to lower levels.

### 2.2 A mathematical model of interacting sub-populations in glioblastoma

Over recent years numerous different approaches have been taken to modelling the growth of GBM; such approaches include examples of multiscale, lattice-based or stochastic models, with each having a focus on capturing particular properties of these complex tumours; a recent comprehensive review of mathematical approaches to modelling GBMs is given by Alfonso et al. in [26]. The approach we follow here, however, is inspired by the “Proliferation-Invasion” (PI) model, which takes the form of the well-known Fisher-KPP equation [27, 28, 29, 30, 31, 32]. The PI model is a minimal model of glioblastoma growth based on the simplified definition of cancer as “uncontrolled proliferation of cells with the ability to invade” [32]; two phenomena that GBMs are well-known to exhibit aggressively. The real power of the PI model lies in its simplicity as model parameters can be estimated from patient MRI scans, allowing patient-specific predictions to be made and inform clinical practice [33, 34, 27, 35]. For this reason, we choose to adopt a similar minimal approach to modelling the co-evolution of EGFR and PDGFRA amplified sub-populations in GBMs.

Our model, therefore, takes the form of an extended PI model to account for the evolution of three genetically-distinct sub-populations defined as tumour cells with the genes encoding for EGFR (*E*), PDGFRA (*P*) and neither (*N*) protein amplified. The model is then given by the system of reaction-diffusion equations:

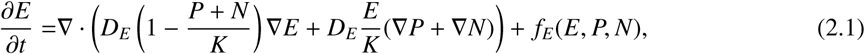

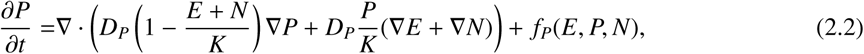

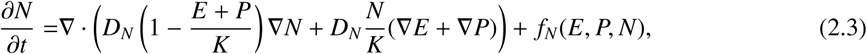

with the initial conditions given by,

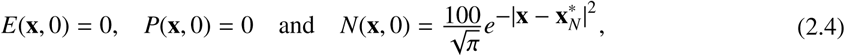

where *E, P* and *N* are the concentrations of each of the tumour cell sub-populations (cells/mm^3^) and 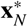 defines the centre of the initial distribution of type *N* tumour cells. Thus we initiate our simulations with no *E* or *P* cells present and 100 cells of type *N*.

Similarly to the PI model, our model essentially consists of two terms to model the evolution of each cell sub-population. The first of these models the ability of each cell type to invade, given by the first term on the right hand side of each equation. Each of these terms model the net migration of each population as a type of diffusion, where the parameters *D*_*E*_, *D*_*P*_ and *D*_*N*_ represent their diffusion coefficients (mm^2^/year). We choose to incorporate this non-linear form of diffusion in our model, since we expect the migration of tumour cell sub-populations to be affected by the presence of other tumour cells and derive these terms following the volume-filling approach of Painter and Hillen [36]. This approach is based on the assumption that only a finite number of cells, the *maximum carrying capacity* (*K*), can fill a given volume, which we assume to be the same for each of the cell populations *E, P* and *N*; other approaches to modelling the migration of multiple GBM tumour cell sub-populations can be found in [37] and [38]. We derive an approximate value for *K* (as in [38]) by assuming that all sub-populations have cells of the same size with a radius of 10µm, yielding a volume of approximately 4.189 × 10^3^µm^3^. Thus, we have a maximum carrying capacity of

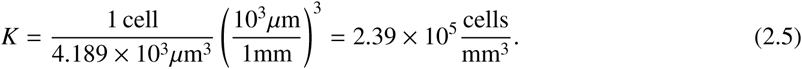

Meanwhile, the uncontrolled proliferation of each tumour cell sub-population is modelled by the terms *f*_*E*_, *f*_*P*_ and *f*_*N*_, which take the form:

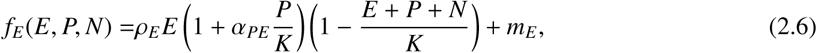

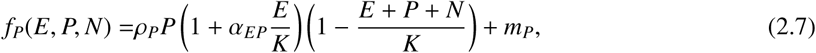

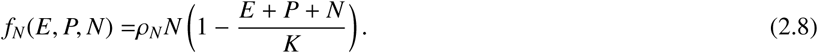

Each of these expressions consists of a term to model the net proliferation of each sub-population. These proliferative terms include a joint logistic growth factor, where we assume there are plenty of other resources such as oxygen and nutrients available and the proliferation of each population is limited only by the availability of space, such that no net proliferation occurs once the carrying capacity, *K*, is reached.

The other factors in the above proliferation terms can be thought of as modified net proliferation rates, where the parameters *ρ*_*E*_, *ρ*_*P*_ and *ρ*_*N*_ represent the intrinsic net proliferation rates of each population (1/year). The parameters *α*_*EP*_ and *α*_*PE*_ measure the effect of sub-population *E* on sub-population *P* and vice versa. For example, if *α*_*EP*_ > 0 in Eq. (2.7), then the presence of the EGFR amplified (EGFRamp) population, *E*, promotes proliferation of the PDGFRA amplified (PDGFRAamp) population, *P*; this could be due to secretion of a growth factor that PDGFRAamp cells are sensitive to, for example. Alternatively, if *α*_*EP*_ < 0, then the net proliferation of PDGFRAamp cells reduces as the density of the EGFRamp population increases and the PDGFRAamp cells are negatively affected. Furthermore, if *α*_*EP*_ = 0, then sub-population *E* has no effect on the net proliferation of *P*. The parameter *α*_*PE*_ is defined analogously and we note that if both *α*_*EP*_ and *α*_*PE*_ are zero, then there are no additional interactions between the two populations, only competition for space. We define the types of interactions that can occur between EGFR and PDGFR amplified sub-populations in our model and summarise them in Table 1. However, in this paper, we will consider just the three types of interactions between EGFRamp and PDGFRAamp cells highlighted in blue, namely neutralism, competition and cooperation.

**Table 1:**
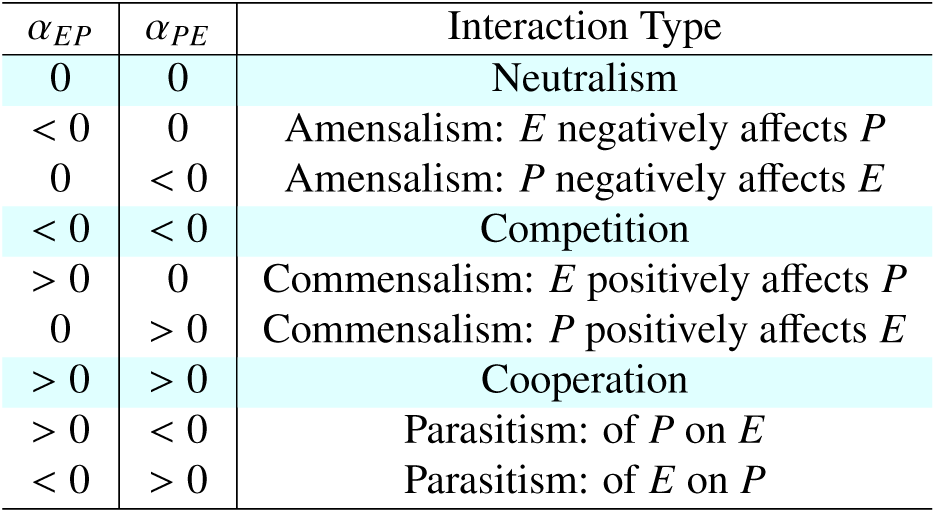
The definitions of interactions that can occur between sub-populations *E* and *P* in our mathematical model as determined by the signs of *α*_*EP*_ and *α*_*PE*_.

| $\alpha_{EP}$ | $\alpha_{PE}$ | Interaction Type |
| --- | --- | --- |
| 0 | 0 | Neutralism |
| < 0 | 0 | Amensalism: $E$ negatively affects $P$ |
| 0 | < 0 | Amensalism: $P$ negatively affects $E$ |
| < 0 | < 0 | Competition |
| > 0 | 0 | Commensalism: $E$ positively affects $P$ |
| 0 | > 0 | Commensalism: $P$ positively affects $E$ |
| > 0 | > 0 | Cooperation |
| > 0 | < 0 | Parasitism: of $P$ on $E$ |
| < 0 | > 0 | Parasitism: of $E$ on $P$ |

Finally, we observe that neither sub-population *E* or *P* is present initially in our model with the initial conditions defined by equation (2.4). These two sub-populations arise in the model through the terms *m*_*E*_ and *m*_*P*_ in Eq.s (2.6) and (2.7); these terms represent the introduction of EGFR and PDGFRA amplified cell populations into the growing tumour that occur due to cells with neither gene amplified, the *N* population, mutating to acquire amplification of each of these genes, respectively. While these mutation events lead to the creation of a single EGFR or PDGFRA amplified cell and there are likely to be many such events occurring during the growth of a GBM, we assume that each of the EGFR and PDGFRA amplified sub-populations only become established within the tumour at most once. Furthermore, since we are using a PDE model more suited to modelling events on the macroscopic scale rather than single cell events, we account for these successful mutation events by introducing a small population of the mutated cells as a distribution. This means that the mutated cell population actually started growing a small amount of time before being introduced in our model, however we assume that this will not have affected the other tumour cell populations present and they only begin interacting and competing for space and resources once the population is of a certain size. We, therefore, choose *m*_*E*_ = *m*_*E*_(**x**, *t*) and *m*_*P*_ = *m*_*P*_(**x**, *t*) to be of the following form,

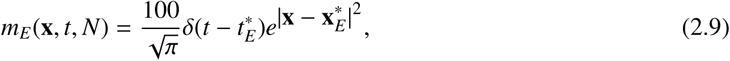

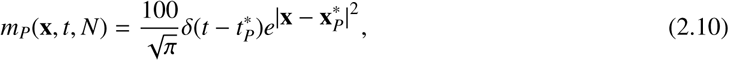

where δ(·) is the Dirac delta function, 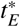 and 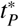 are the times at which populations of 100 EGFR and PDGFRA amplified cells, *E* and *P*, are introduced as Gaussian distributions centred at 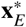 and 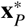, respectively. We note that, since we are working with a continuum model, we are required to introduce the cells as a distribution and also choose the size of this distribution; there are several possible choices for this and we found that the choice of distribution did not qualitatively change our model simulations, as long as they were chosen consistently, due to the smoothing effects of diffusion in the model.

The times, 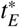 and 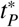, and locations, 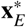 and 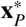, that these distributions of mutated populations are introduced into our simulations, however, do have an effect on our results. This will be illustrated and discussed in the next section, where we explore the behaviour of our model under different interaction assumptions on a one-dimensional tumour slice with according assumptions about symmetry and compare the simulation outputs to patient biopsy data. All model simulations were produced using MatLab R2017a to implement a finite difference scheme with zero flux boundary conditions and using a large enough spatial domain such that tumour growth is far away from the boundaries to avoid introducing any boundary condition artefacts. In our finite difference scheme a spatial mesh size of 0.25mm and a time step of 1/1500 years were used.

Before moving on to presenting the results of this paper, we observe briefly that under the assumption that the EGFR and PDGFRA amplified sub-populations do not interact with one another, i.e. *α*_*EP*_ = *α*_*PE*_ = 0, and all three sub-populations diffuse and proliferate at the same rates, i.e. *D*_*E*_ = *D*_*P*_ = *D*_*N*_ and *ρ*_*E*_ = *ρ*_*P*_ = *ρ*_*N*_, then the system can be reduced to a single equation governing the total population of tumour cells, *T* = *E* + *P* + *N*, for all 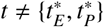; this equation is the well-known, clinically significant Proliferation-Invasion (PI) model that describes the evolution of a single homogeneous population of GBM tumour cells mentioned at the start of this section. Therefore, we note that our model is consistent with the PI model when used to model a single homogeneous population *T*.

## 3 Results

### 3.1 Our model predicts that distinct competing sub-populations of tumour cells can coexist in the same tumour region

Intuitively, we expect that cell populations actively competing with one another may be less likely to coexist within the same region of a tumour and that their coexistence may indicate a cooperative relationship, as Snuderl et al. [21] suggest after finding intermingled sub-populations of EGFR and PDGFRA amplified sub-populations in a small number of GBM samples. In our model, however, we find that EGFR and PDGFRA amplified sub-populations can be found to coexist in areas of the tumour region despite actively competing with one another; we find that any 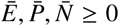 satisfying 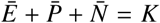 is a spatially homogeneous steady state and can be connected to other spatially homogeneous steady states satisfying the same condition, or the trivial steady state 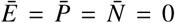, by travelling wave-like solutions expanding outwards from the origin of the tumour. Indeed, such co-occurrence of EGFR and PDGFRA amplified cell sub-populations can be observed with competitive, cooperative or neutral interactions; an example of simulations with different *α*_*EP*_ and *α*_*PE*_ values to represent each of these interaction types is shown in Fig. 1, where such co-occurrence is observed.

**Figure 1:**
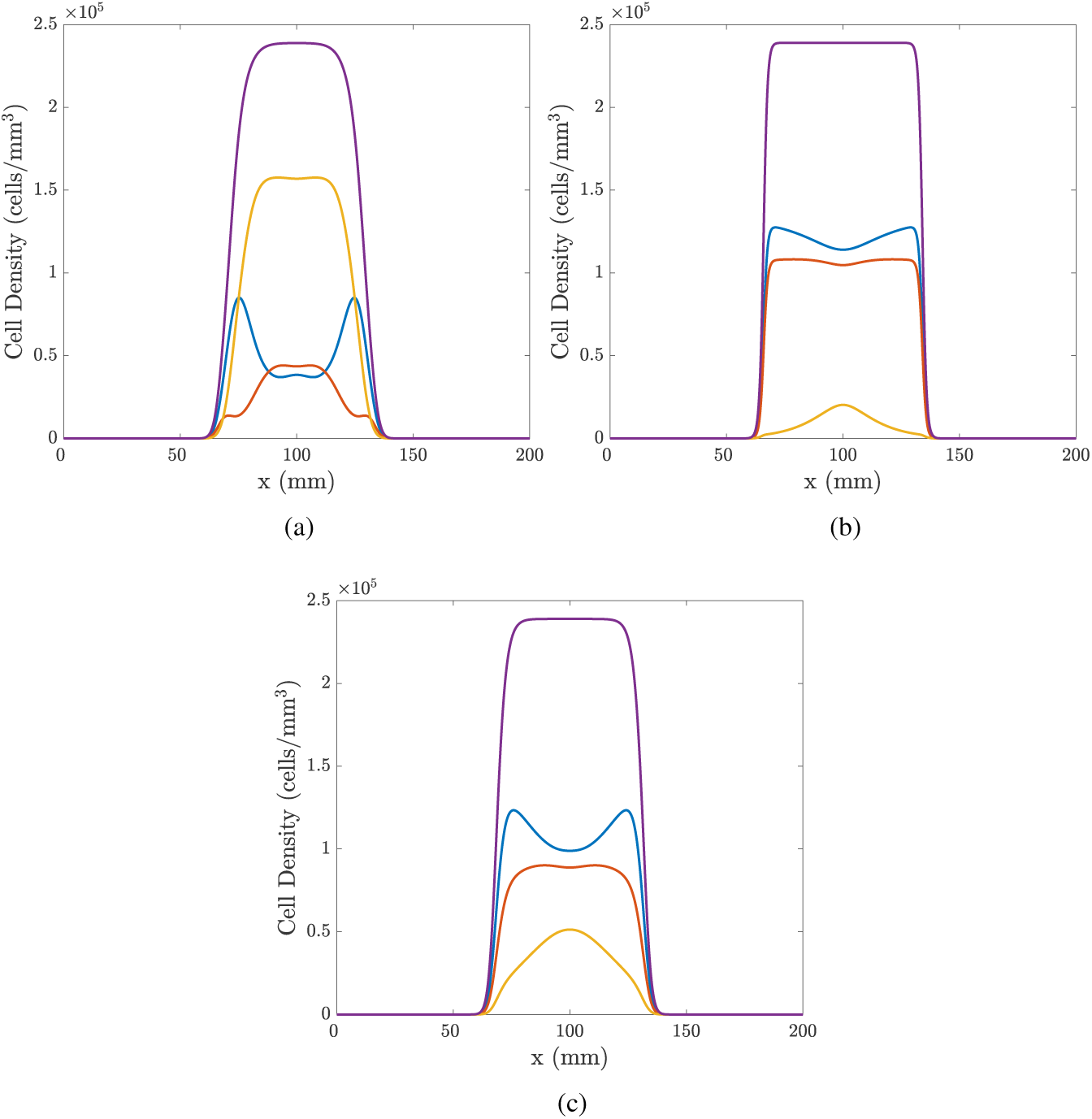
Simulations in 1D of the model given by Eq.s (2.1)-(2.8) with parameters: *ρ*_*E*_ = 35.4, *ρ*_*P*_ = 33 and *ρ*_*N*_ = 30 1/year; *D*_*E*_ = *D*_*P*_ = *D*_*N*_ = 30mm^2^/year;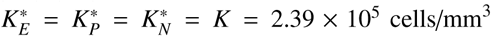; 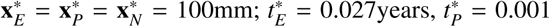years. The interactions in each simulation are chosen to be (a) competition, *α*_*EP*_ = *α*_*PE*_ = −5, (b) cooperation, *α*_*EP*_ = *α*_*PE*_ = 5 and (c) neutralism, *α*_*EP*_ = *α*_*PE*_ = 0. Population *E* is shown in blue, *P* in red, *N* in yellow and the total population, *T* = *E* + *P* + *N*, in purple.

While this demonstrates that the coexistence of EGFR and PDGFRA amplified cell populations in one known region of a tumour can occur when they are competing, cooperating and evolving neutrally, it highlights the need for more information to determine the nature of such interactions between co-occurring cells *in vivo*. In this work we hope to shed some light on this by studying amplification patterns observed across image-localized biopsies from a cohort of patients and the patterns that emerge in simulations of our model, given by Eq.s (2.1)-(2.10), under different interaction assumptions.

### 3.2 Biopsy data

A total of 120 biopsies were collected from 25 GBM patients, with 2-14 collected from each individual. Of these biopsies, 95 samples from 25 patients contained adequate tumour and/or DNA content for EGFR and PDGFRA amplification status to be determined successfully through aCGH analysis. EGFR amplification was the more commonly observed genetic alteration, with 73/95 samples having a CNA value associated with EGFR amplification, whereas 28/95 were determined to be PDGFRA amplified. Of these amplified samples, 22 were found to have amplification of both the EGFR and PDGFRA genes. For each patient, we then determined the proportion of their biopsies that were found to be amplified in neither gene, only the EGFR gene, only the PDGFRA gene and, finally, both of the EGFR and PDGFRA genes simultaneously. The proportions calculated for each of the 25 patients are summarised as a box plot in Fig. 2(a) and the mean of these proportions are shown as a spider plot in Fig. 2(b).

**Figure 2:**
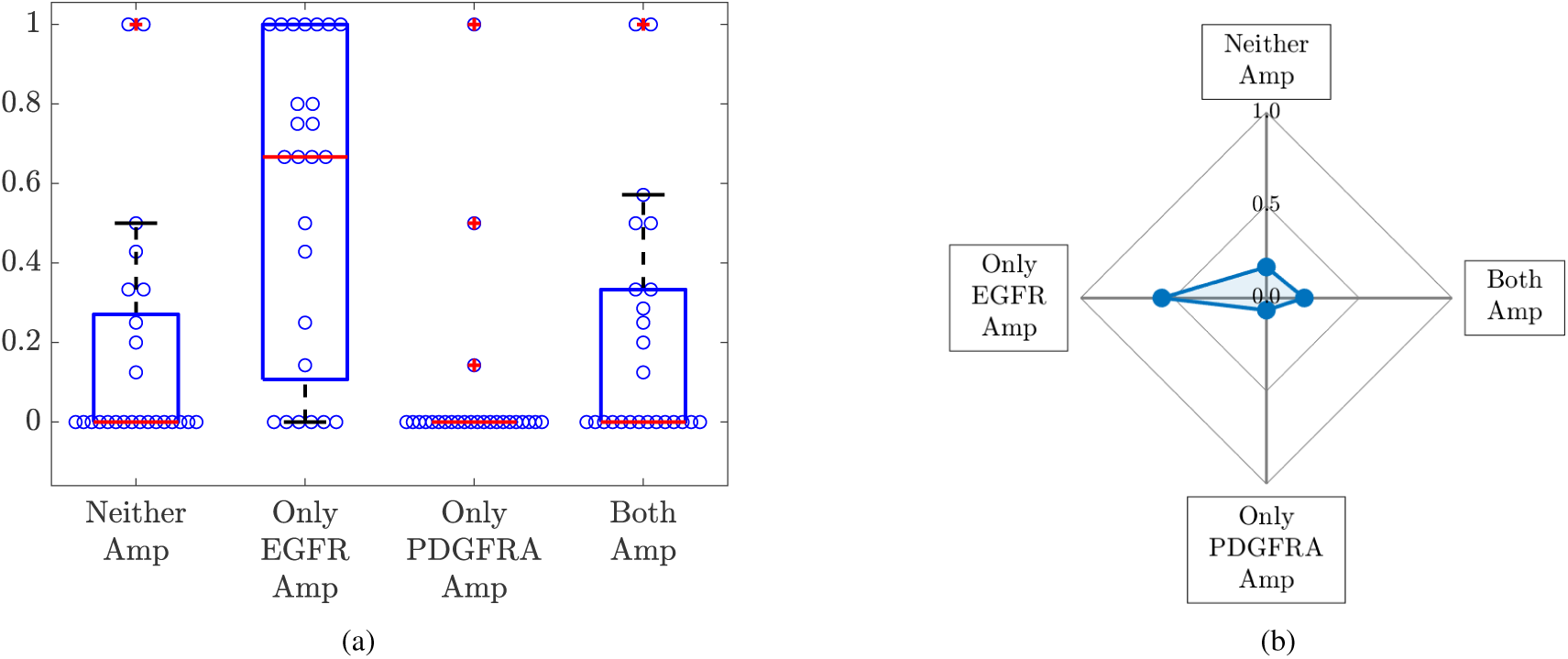
(a) A box plot summarising the proportion of each individual’s biopsies that were determined to be amplified in neither gene (Neither), only the EGFR gene (Only EGFR Amp), only the PDGFRA gene (Only PDGFRA Amp) and both of the EGFR and PDGFRA genes (Both Amp) for the 25 patients. Each of the blue circles overlaid on the box plot represents the relevant proportion of an individual’s biopsies for each category. The means of these proportions across the 25 patients are shown in (b).

### 3.3 Simulation results

In order to test whether neutral, competitive or cooperative interactions between EGFR and PDGFRA amplified sub-populations best describe the patterns observed in the biopsy data, we run numerical simulations of the model described in Section 2.2 and compare the outputs to the mean proportions of biopsies amplified in neither gene, only the EGFR gene, only the PDGFRA gene and both of the EGFR and PDGFRA genes shown in Fig. 2(b).

To do this, we begin by assuming that amplification of the EGFR or PDGFRA gene does not result in either sub-population, *E* or *P* in the model, acquiring any selective advantages over non-amplified cells. In other words, we choose all proliferation and invasion parameters to be the same for each population, i.e. *ρ* = *ρ*_*E*_ = *ρ*_*P*_ = *ρ*_*N*_ and *D* = *D*_*E*_ = *D*_*P*_ = *D*_*N*_. Since the biopsy data presented in Fig. 2(b) is the mean of a cohort containing 25 individual tumours, which will vary in their invasive (quantified by *ρ*/*D* in the PI model [33]) and proliferative (quantified by *ρ*) qualities, we choose to simulate using a variety of *ρ* and *D* pairs to reflect the heterogeneity seen in the patient cohort. We therefore produce four simulations using two values for each of *ρ* and *D* to mirror the range of parameters observed in unpublished patient databases and assume this cohort are similarly distributed; to represent high parameter values we use *ρ* = 30/year and *D* = 30mm^2^/year and for low values we use *ρ* = 3/year and *D* = 3mm^2^/year as in [39]. We also assume that any interactions occurring between the EGFR and PDGFRA amplified cells affect each sub-population to the same degree, i.e. *α* = *α*_*EP*_ = *α*_*PE*_. To represent the two cases of competition and cooperation, we simulate with *α* = −5 and 5, respectively, while the neutralism case is *α* = 0. Simulations are then produced using these sets of parameter values on the spatial domain **x** = [0, 200] mm, which is large enough to avoid introducing boundary artefacts given that we only run simulations to a biologically relevant size and all sub-populations, *E, P* and *N*, are introduced at the centre of this domain 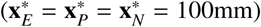, with *N* at time *t* = 0 and *E* and *P* at the later times 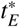 and 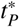, respectively. For now we assume that the EGFR and PDGFRA mutation events that lead to the introduction of the EGFR and PDGFRA amplified sub-populations, *E* and *P*, in our model occur at the same time, so we take 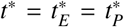. Since the time evolution of our simulated tumours scales with *ρ*, rather than introduce these populations at a fixed point in time, we introduce them after a given number of proliferation events have occurred, i.e. a fixed point in the evolution of the tumours. In this case we choose *t*\* to be the first time point after the tumour has grown to contain a total of 600 cells of type *N*. We note that, while this may seem particularly early in the evolution of our simulated tumours, it is necessary in order to be able to investigate the amplification patterns we observe under different interactions, since if we introduce the EGFR and PDGFRA amplified populations much later we do not see them growing to detectable levels in our simulations.

Finally, we choose a biologically relevant size to represent a typical size of a GBM tumour at the time of diagnosis, shortly after which patients will usually undergo surgery to remove as much of the tumour as feasible, with biopsy samples being collected at this time. One way to measure the typical size of a GBM at diagnosis is to segment the tumour volume visible on a T1-weighted MRI with gadolinium contrast (T1Gd MRI), a type of scan that shows the most dense area of the tumour and is typically used in the process of diagnosing a patient with a GBM. From this, the diameter of the volume-equivalent sphere can then be computed and used as a measure to indicate the size of the tumour lesion, with the average diameter at diagnosis being 36.2mm in unpublished data. Following Swanson et al. [27], we relate the tumour volume visible on a T1Gd MRI to the volume of tumour that has a tumour cell density greater than 80% of the carrying capacity. Therefore, we choose to run simulations until the width of the total tumour cell population, *T* = *E* + *P* + *N*, above the 0.8*K* threshold is 36.2mm and use this as a proxy for the size of a tumour at the time of diagnosis.

We then run the numerical simulations until they reach this biologically relevant size with the parameter sets described above. Since our patient data looks at the proportions of biopsies containing EGFR and PDGFRA amplified cells above a given density threshold, which we chose to be 10% of the tissue sample, we treat each spatial mesh point in our simulations as if it were a biopsy point. Thus, we calculate the proportions of mesh points with neither gene, only the EGFR gene, only the PDGFRA gene and both genes amplified, by first counting the number of mesh points with *T* = *E* + *P* + *N* > 0.1*K*, as illustrated schematically in Fig. 3(a), and then counting the mesh points with the relevant sub-populations above and below this threshold. So, for example, the proportion of mesh points with only EGFR amplified, *A*_*E*_, is calculated as

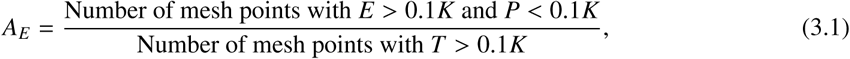

and the proportions with neither gene, only the PDGFRA gene and both genes amplified, *A*_*N*_, *A*_*P*_ and *A*_*B*_, are calculated in a similar way. The mean proportions from the simulations run using each of the four *ρ* and *D* pairs with the different cases of competition, cooperation and neutralism are shown in Fig. 3(b). From this plot, we see that all simulations are classed as neither gene amplified in the competitive case and this proportion decreases as we move through the neutralism case to the cooperative case and the proportion with both genes amplified increases, which is as we would intuitively expect to see. We notice that in all three cases the proportions of simulations with only one of the genes amplified are zero; again, this is expected as populations *E* and *P* have the same proliferation and invasion parameters (*ρ* and *D*) and are both introduced at the same time and place and, thus, are effectively the same so we expect to see them together (or not at all in the competitive case).

**Figure 3:**
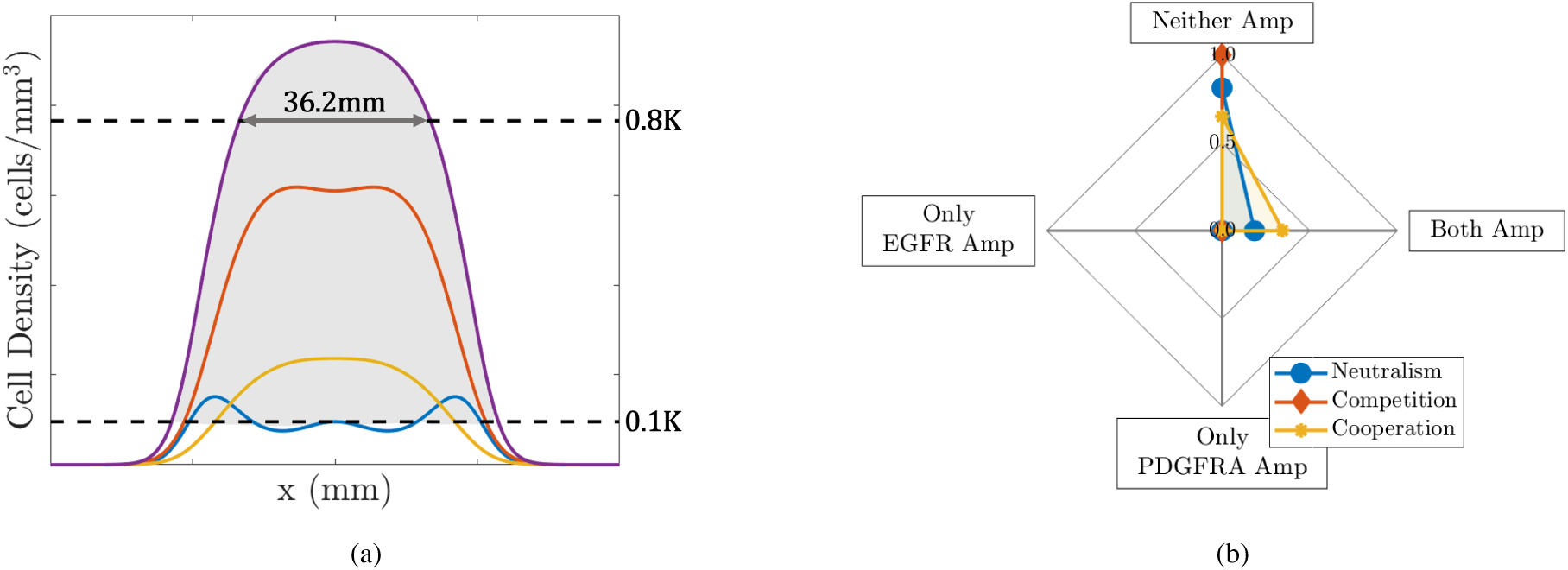
(a) Schematic illustrating the biologically relevant size that tumours are simulated to and area (shaded in grey) indicating the spatial mesh points with the total tumour cell population (purple curve) above the threshold of 10% of the carrying capacity. Other curves represent the individual tumour cell populations; *E* (blue), *P* (red) and *N* (yellow). (b) Plot showing the mean proportions of simulations with neither gene (Neither Amp), only the EGFR gene (Only EGFR Amp), only the PDGFRA gene (Only PDGFRA Amp) and both genes (Both Amp) amplified under different interactions when we assume that the sub-populations are dynamically the same, i.e. all populations have the same proliferation and invasion parameters, *ρ* = *ρ*_*E*_ = *ρ*_*P*_ = *ρ*_*N*_ and *D* = *D*_*E*_ = *D*_*P*_ = *D*_*N*_, and *E* and *P* populations are introduced at the same position and time, 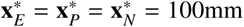 and 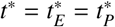, as described in Section 3.3.

Clearly, the patterns of neither, only EGFR, only PDGFRA and both amplified proportions we see in these simulations do not reflect the patterns of amplification we see in the biopsy data in Fig. 2(b); one obvious difference is the proportion of only EGFR amplified biopsies, which is above 0.5 in the data and is 0 in the simulations shown in Fig. 3(b). Since EGFR and PDGFRA amplified cells are not exclusively found in the same biopsies in the patient data and the proportions of biopsies with only one gene amplified also differ, this indicates the two populations must differ in their dynamics in some way. There are several possible ways that differences between the EGFR and PDGFRA amplified sub-populations can occur in our model: by giving them a selective advantage (changing *ρ*_*E*_, *ρ*_*P*_, *D*_*E*_ or *D*_*P*_); by changing the phylogenetic ordering of mutations (changing 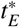 or 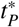); by changing the location that mutations arise in the evolving tumour (changing 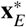 or 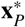); by changing the strength of interaction felt by each population (changing *α*_*EP*_ or *α*_*PE*_); or, finally, by changing any combination of these factors. Since little is known about the type of interactions occurring between these amplified cell sub-populations, we continue to assume that the effect is symmetric and they are affected by the presence of each other to the same degree. Therefore, in the following sections of this paper, we begin to explore the effect of introducing selection advantages, changing the phylogenetic ordering of mutations and the location that the amplified populations are introduced.

#### 3.3.1 Selection advantages

The amplification of EGFR and PDGFRA genes are often considered to be among the key mutations driving oncogenesis and tumour growth [21, 22]. Since these genes are both members of the RTK family of cell surface receptors that play an important role in the regulation of cell proliferation, metabolism and survival [12] and tumours identified to be EGFR amplified have shown to be more invasive [6, 15], it is reasonable to consider that amplification of these genes may drive the growth of tumours through increasing the intrinsic proliferative and invasive abilities of these cell sub-populations. We can explore the effects of this by giving the EGFR and PDGFRA amplified sub-populations in our model different selection advantages through changing the appropriate parameters, namely *ρ*_*E*_, *ρ*_*P*_, *D*_*E*_ and *D*_*P*_.

As we see a much higher proportion of biopsies with only EGFR amplified in the patient data (Fig. 2(b)), this could indicate that EGFR amplified cells have a selective advantage over the PDGFRA amplified cells and those with neither gene amplified. Thus, we explore the effects of giving EGFR amplified cells invasive and proliferative advantages; in Fig. 4, plots are produced from simulation results in the same way as previously described, but with (a) a 50% invasive advantage for the EGFR amplified cells and (b) additionally with a proliferative advantage. In both of these cases, there are now large proportions of simulations with EGFR amplified and without PDGFRA amplified cells present, particularly in the competitive and neutral cases. In both competitive cases in Fig.s 4(a) and 4(b), we observe nowhere where both of the EGFR and PDGFRA genes are found to be amplified, unlike the patient data in Fig. 2(b) where this was approximately 20% of biopsies. Meanwhile, the corresponding cooperative cases both gave the highest level of points with both genes amplified and the lowest levels with only EGFR amplified. In Fig. 4, we also explored the effects of affording the PDGFRA amplified sub-population a (c) 50% invasive and (d) 50% proliferative advantage over non-amplified cells while giving EGFR amplified cells the same advantages as in (b). These resulted in qualitatively similar amplification patterns to those observed in (b) and (a), respectively, with the exception of the competitive case in (d) where a small proportion of simulations with only PDGFRA amplified were observed. Further plots exploring the effects of selection advantages can be found in the supplementary material in Fig. S1 and are not presented here for brevity. However, while we find that giving either, or both, of the amplified populations invasive and proliferative advantages over non-amplified cells improves the amplification patterns we see in simulations with respect to the biopsy data, it is not a perfect fit. Investigating other degrees of invasion and proliferation advantages may improve the results we see, since we only looked at quite large advantages of 50% of the respective parameters, which is something we may investigate in future work. Here, however, we move on to explore the effects that the timing of mutations have on the results we see.

**Figure 4:**
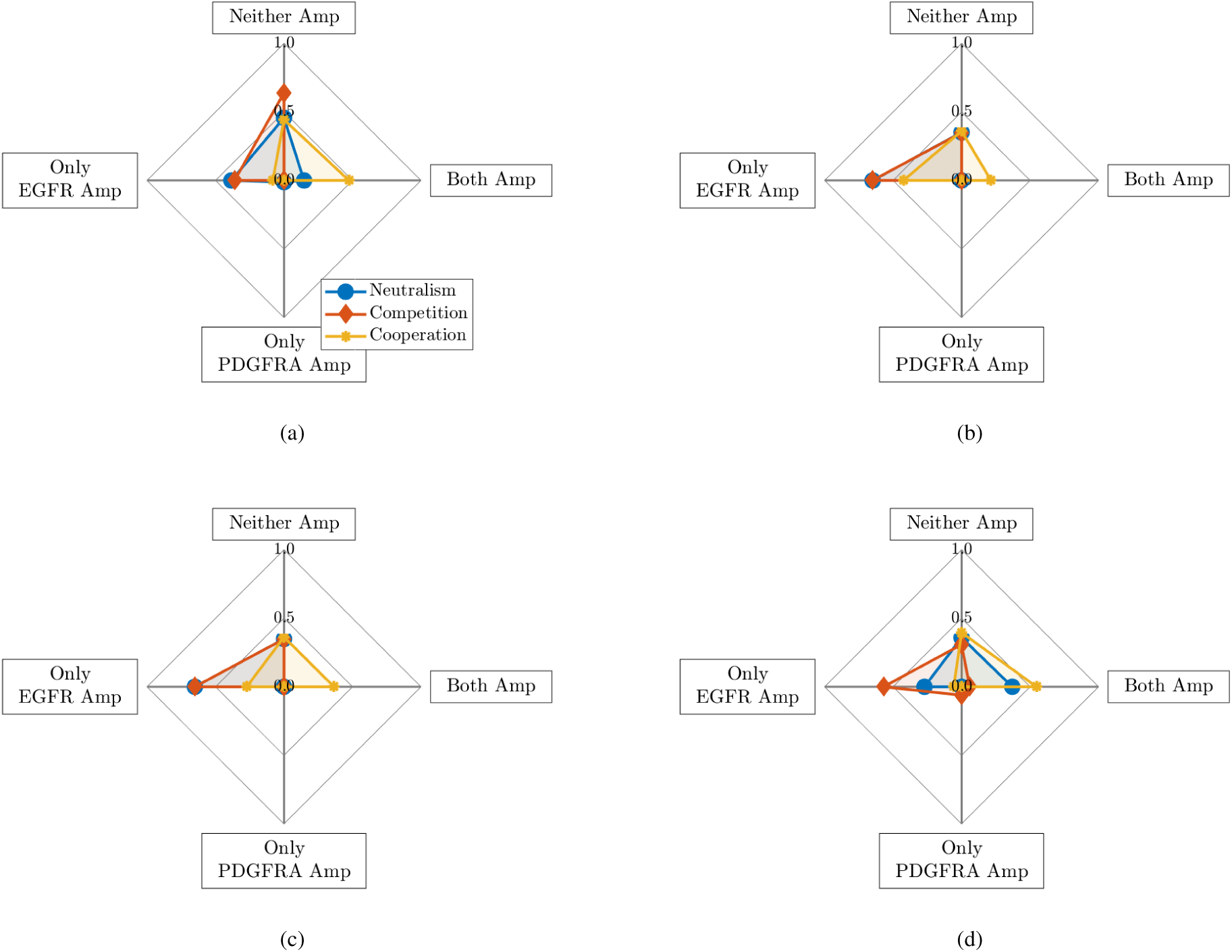
Plot showing the mean proportions of simulations with neither gene (Neither Amp), only the EGFR gene (Only EGFR Amp), only the PDGFRA gene (Only PDGFRA Amp) and both genes (Both Amp) amplified under different interactions when the *E* and *P* sub-populations are given various selection advantages: (a) EGFR 50% invasive advantage (*D*_*E*_ = 1.5*D*_*P*_ = 1.5*D*_*N*_); (b) EGFR 50% proliferative and invasive advantage (*ρ*_*E*_ = 1.5*ρ*_*P*_ = 1.5*ρ*_*N*_ and *D*_*E*_ = 1.5*D*_*P*_ = 1.5*D*_*N*_); (c) EGFR 50% proliferative and invasive advantage, PDGFRA 50% invasive advantage (*ρ*_*E*_ = 1.5*ρ*_*P*_ = 1.5*ρ*_*N*_ and *D*_*E*_ = *D*_*P*_ = 1.5*D*_*N*_); (d) EGFR 50% proliferative and invasive advantage, PDGFRA 50% proliferative advantage (*ρ*_*E*_ = *ρ*_*P*_ = 1.5*ρ*_*N*_ and *D*_*E*_ = 1.5*D*_*P*_ = 1.5*D*_*N*_)

#### 3.3.2 The phylogenetic ordering of mutations

In all simulations presented up until this point, we have assumed that the mutations leading to the establishment of EGFR and PDGFRA amplified cell sub-populations occur at the same time. While this could be the case, a study reconstructing the phylogeny of GBMs identified EGFR and PDGFRA amplification as early and late events during tumour progression [8]. From an analysis of multiple spatially distinct samples from 11 GBMs, it was inferred that alterations that were more common occurred earlier in the evolution of the tumour compared to those only present in a smaller subset of cells. Alterations in gene copy numbers on the chromosomes where the EGFR and PDGFRA genes are found tended to occur in the early and middle phases of tumour growth, respectively [8]. Therefore, this timing, or *phylogenetic ordering*, of mutations may be affecting the proportions of EGFR and PDGFRA amplified biopsies across the patient cohort (Fig. 2(b)) and so we undertake a brief exploration of the effect it has in our simulations.

Therefore, we assume once more that all cell sub-populations have the same proliferation and invasion parameters, i.e. *ρ* = *ρ*_*E*_ = *ρ*_*P*_ = *ρ*_*N*_ and *D* = *D*_*E*_ = *D*_*P*_ = *D*_*N*_ and, as described before, we simulate using four different *ρ* and *D* pairs and the same assumptions described in Section 3.3, with the exception of changing the times that the EGFR (*E*) and PDGFRA (*P*) amplified populations are introduced, i.e. 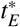 and 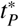. As before, instead of choosing a fixed time at which to introduce these sub-populations in our model simulations, we choose this time based on the number of cells that the evolving tumour has grown to. In the previous simulations, both populations were introduced after the non-amplified (*N*) population had grown to a size of 600 cells and so we now choose to investigate the patterns of gene amplification we see when the *E* and *P* populations are introduced earlier and later than this. Thus, we define a vector of possible introduction times, 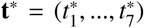 as follows: 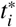 is defined as the first time point in our simulations after the growing *N* population has reached a total of 200 + 100*i* cells, for *i* = 1, …, 7. Implementing our model with 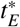 and 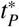 taking each of these values (a total of 49 simulations), we gain some insight into the effect that changing the time of mutations has on our simulations.

Again, for brevity we do not present all the simulation results here and the remainder can be found in the supplementary material (Fig.s S2 and S3), however a set of results are shown here in Fig.5. This figure shows the average amplified proportions in seven sets of simulations under neutral, competitive and cooperative interaction assumptions, where cells of type *P* are introduced at 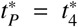 in each set, but the *E* population is introduced at each of the seven possible times given by the vector *t*\*. The points on the graph where 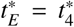 (marked by the dotted vertical line) are the same data represented in Fig. 3(b) as the EGFR and PDGFRA amplified populations are introduced at the same time. Since evidence suggests that EGFR amplification is an earlier event than PDGFRA amplification, of particular interest is to the left of this dotted line on the graph, where we find that the proportion of simulations with neither gene amplified decreases monotonically in all three interaction cases and the proportion with only EGFR amplified increases monotonically; the only exception to this being the cooperative case, where much higher proportions of points with both genes amplified are seen. Meanwhile, as *E* cells are introduced later than *P* cells, we see the proportions changing as we would expect; the only PDGFRA amplified proportion increasing, both amplified proportions decreasing or remaining at zero and the non-amplified proportion either increasing or remaining the same. We note the slight decrease in the proportion of neither amplified cells when the EGFR amplified population is introduced at 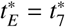 in the competitive case, this is because the PDGFRA population is allowed enough time to proliferate to a size where the introduction and competitive interactions of EGFR amplified cells has a smaller relative effect on their proliferation. As with the previous section where we looked at the effect of giving the amplified sub-populations selection advantages, we find that changing the timing of introducing the mutated populations in our simulations does not fit the biopsy data perfectly, but has some of the desired effects. Such as, increasing the proportions of simulations with only EGFR amplified and decreasing those with neither gene amplified when the introduction of the EGFR amplified population is assumed to be an earlier event in tumour evolution.

**Figure 5:**
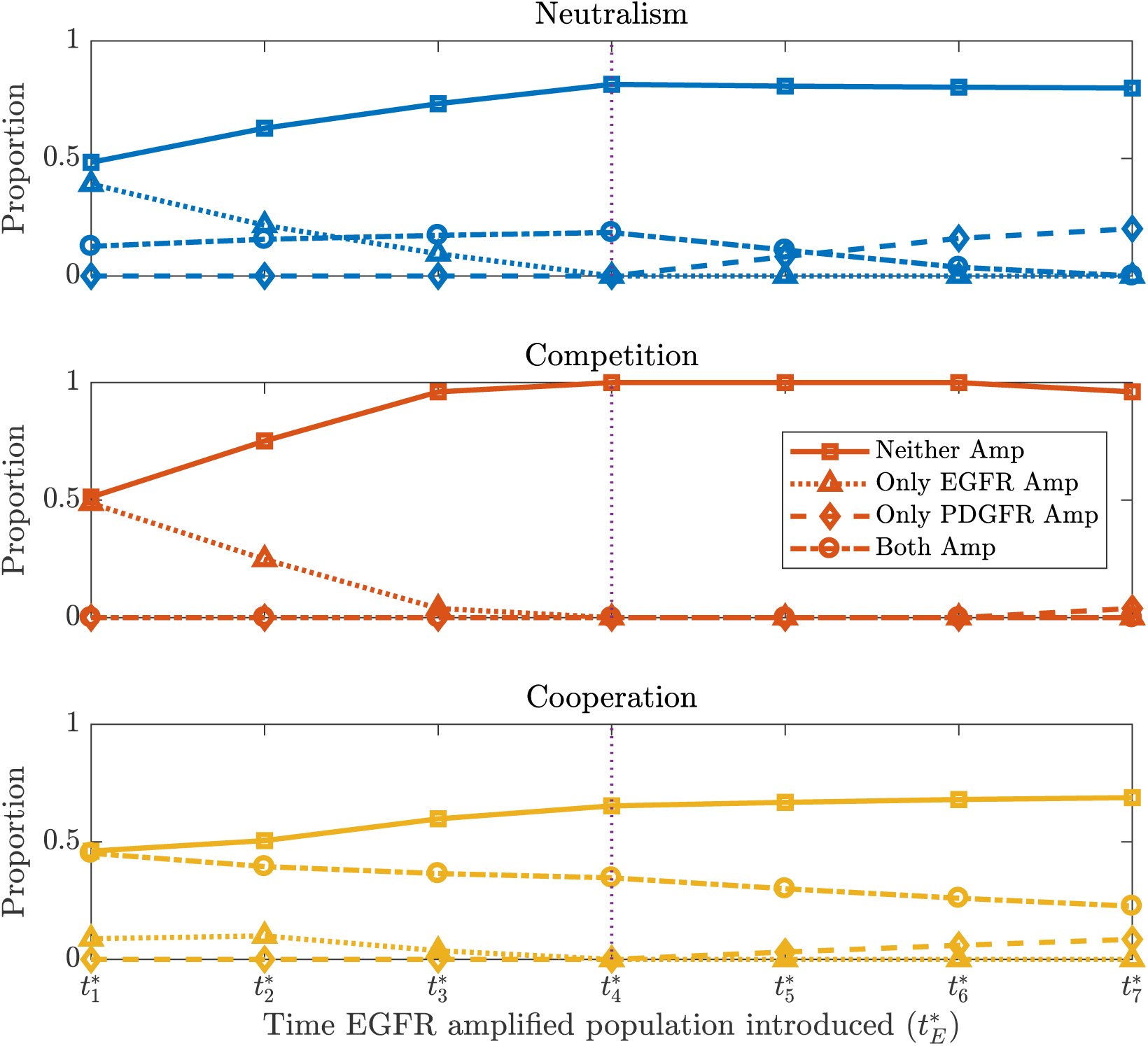
Plots showing the mean proportions of simulations with neither gene (Neither Amp), only the EGFR gene (Only EGFR Amp), only the PDGFRA gene (Only PDGFRA Amp) and both genes (Both Amp) amplified as 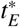 is varied, under neutral, competitive and cooperative interactions. The PDGFRA amplified population is introduced at the fixed time 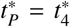 (denoted by the vertical dotted line) and other parameters and assumptions are as described in Section 3.3.

#### 3.3.3 The location of mutations

Another factor that could result in some biopsies having only EGFR and others only PDGFRA amplified is that the mutations occurred in different places, resulting in the populations occupying different, spatially separated regions of the tumour. In all previous simulations presented in this paper we assumed that the mutation events leading to the establishment of EGFR and PDGFRA amplified sup-populations in our model occurred in the center of the growing tumour; this was a reasonable place to explore the effects of selection advantages and timing of mutations from, since it is where most proliferation is taking place in our model in the early phases of tumour growth and, therefore, where we may expect more mutations to appear. However, cells at the centre of the tumour also experience more competition for space, a growth limiting resource in our model, as this is where the highest tumour cell density is in these early growth phases. Therefore, we now explore the effects of introducing the EGFR and PDGFRA amplified sub-populations away from the centre of the tumour where there is less competition for space. Thus, we define a vector of possible introduction locations, 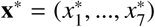 as follows: 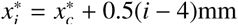, where 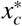 is the mesh point at the centre of the tumour. In this way, 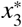 is the mesh point lying 0.5mm to the left of the tumour centre and 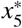 is 0.5mm to the right. We produce simulations using the same parameters and assumptions described in Section 3.3, but with 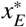 and 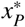 each taking the values given by the vector **x**\*, so a total of 49 simulations. As in previous sections, we do not present all simulation results here and refer the reader to Fig.s S4 and S5 in the supplementary material for more results. We do however show two simulation results in Fig. 6 showing the proportions when EGFR and PDGFRA amplified populations are introduced (a) 0.5mm and (b) 1mm to the left and right of the tumour center, respectively. In both of these cases we see distinct regions with EGFR and PDGFRA amplification forming in the simulated tumours and equal proportions with only one gene amplified. As found in the previous two sections of this paper, only changing where the amplified sub-populations are introduced does not fully explain the patterns of amplification we observe in the biopsy data, however it does produce some of the desired effects, such as increasing the proportion of simulations with only one of each gene amplified.

**Figure 6:**
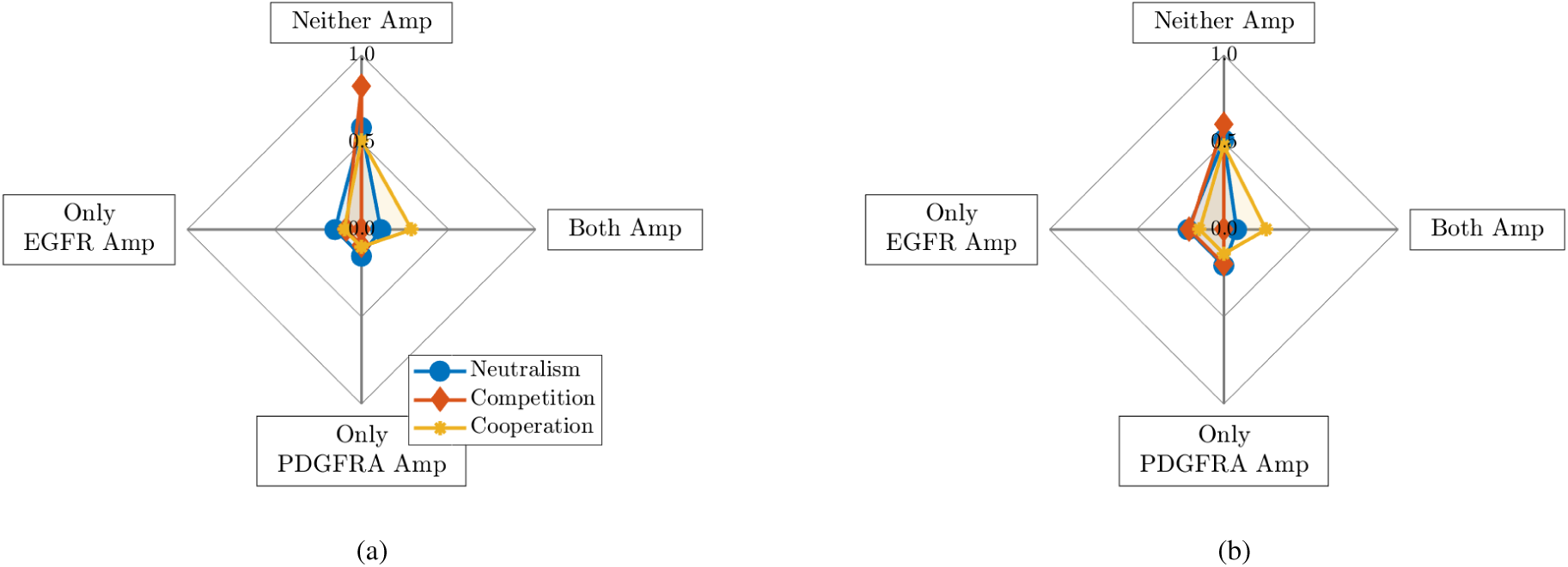
Plots showing mean proportions of simulations with neither gene (Neither Amp), only the EGFR gene (Only EGFR Amp), only the PDGFRA gene (Only PDGFRA Amp) and both genes (Both Amp) amplified when EGFR and PDGFRA amplified sub-populations are introduced (a) 0.5mm to left and right of center (i.e. 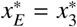 and 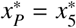) and (b) 1mm to left right of center (i.e. 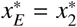 and 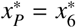), respectively.

## 4 Discussion

In this paper, we have presented a novel mathematical model of the co-evolution of three distinct tumour cell sub-populations to investigate the nature of interactions between cells with two common mutations occurring in GBMs, namely amplification of the genes encoding the EGFR and PDGFRA proteins. We have used a PDE-based formalism, which reduces to the well known PI model [27, 28, 29, 30, 31, 32] if we assume that these genetic differences do not change the phenotype of the cell populations and instead compose a single phenotypically homogeneous population of cancerous cells. We then conducted an *in silico* investigation into the levels of EGFR and PDGFRA amplified sub-populations observed under competitive, cooperative and neutral interactions between these cell types and compared our results to population levels of amplification observed in image-localized biopsy data from a cohort of GBM patients, where a high proportion of biopsies had only the EGFR gene amplified, a smaller proportion had both or neither gene amplified and very few showed amplification of only the PDGFRA gene (see Fig. 2(b)).

In carrying out computational simulations, we found that the amplification patterns observed in simulated tumours under each of the different interaction assumptions did not match those observed across the patient biopsy data when we assumed that cells with the EGFR and PDGFRA genes amplified did not differ in their dynamics, shown in Fig. 3(b). We note that this was to be expected and is consistent with research suggesting dynamical differences between these sub-populations. For example, a study reconstructing the phylogeny of GBMs suggests that EGFR amplification is a mutation that arises earlier than amplification of the PDGFRA gene [8], while other studies have found that tumours with EGFR amplified are more invasive, which could indicate that EGFR amplified cells themselves are more invasive [14, 15]. Following this, we explored the effects of introducing differences between the sub-populations in our simulations through changing various model parameters. Since a high proportion of biopsies across the patient data have the EGFR gene amplified but no amplification of the PDGFRA gene, we investigated the effects of giving EGFR amplified cells various selection advantages over PDGFRA and non-amplified cell types, shown in Fig. 4 and Fig. S1 in the supplementary material. While these simulations do not achieve the same amplification patterns observed in the biopsy data, affording EGFR amplified cells a selection advantage does help to produce the high levels of EGFR amplification required, particularly in the competition and neutral interaction cases. We then chose to investigate the effects of changing the phylogenetic ordering of mutations and found that introducing the EGFR amplified sub-population earlier in the evolution of our simulated tumours also helped to produce the desired higher level of EGFR amplification under competitive and neutral interaction assumptions, whereas the proportion of points with both genes amplified was much higher than observed in the biopsy data in the cooperative case. Finally, we looked at introducing the EGFR and PDGFRA amplified sub-populations in different locations in the growing tumour simulations. We found that changing these locations did not have such a large effect on the amplification patterns observed as the selection advantages and phylogeny did, however it could be an important factor in replicating amplification patterns observed in individual GBMs with distinct regions of EGFR and PDGFRA amplification. Although the simulation results presented in this paper do not perfectly match the patterns of EGFR and PDGFRA amplification observed across the patient biopsy data, our *in silico* modelling approach has allowed us to investigate how different interaction assumptions influence the amplification patterns in simulated tumours and explore the effects of changing the parameters and the timing and position of sub-population introductions in our model; we found that some of these changes improved our simulation results with respect to the biopsy data and were also consistent with suggestions about EGFR amplified sub-populations found in the literature.

In this paper, we only explored each factor individually, whereas a combination of factors relating to the selection advantages of EGFR and PDGFRA amplified cells as well as the timing and location of mutations are likely to be influencing the amplification patterns across the biopsy data. This combination of factors makes it is difficult, at this stage, to deduce the nature of interactions between cell types that is driving these patterns. Furthermore, we only studied one strength of competitive and cooperative interactions between EGFR and PDGFRA amplified sub-populations, whereas these cells may be interacting to stronger or weaker degrees. Both of these are potential limitations of this paper that we would like to address and investigate in future work.

We also note that in this work we have avoided modelling single cell events in a macroscopic setting, by assuming that each of the EGFR and PDGFRA amplified sub-populations only become established within a tumour at most once and introduce a small distribution of cells accordingly. It is possible, however, that multiple mutation events resulting in EGFR and PDGFRA amplification in cells could be occurring in an evolving tumour and it may be more appropriate to model such single cell events in a micro- or meso-scopic setting. However, since we are interested in population level patterns of amplification and are working with large numbers of cells, this would be computationally expensive. In order to better capture single cell events while avoiding a large computational burden, it may be appropriate to consider a hybrid-modelling approach, similar to that described by Smith and Yates [40]. Though in this work, we decided to represent a successful mutation event by introducing a small population of cells in our continuum PDE model and leave such considerations for future work.

Determining the nature of interactions between EGFR and PDGFRA amplified sub-populations in GBMs is a complex biological problem, with factors relating to selection advantages and the phylogeny of these tumours influencing the balance of populations we see in a tumour, as we have demonstrated with our *in silico* investigation in this paper. To be able to untangle the influence from and the nature of interactions from the effects of these other factors, more data may be required. In this study, we had biopsy data for a cohort of 25 patients and a larger number may enable us to gain more insight into the pattern of EGFR and PDGFRA amplification. Furthermore, data at multiple time points, such as biopsies sampled from primary and recurrent tumours, would enable us to extract more dynamic information about the co-evolution of these genetically-distinct sub-populations; this presents challenges, however, as treatment effects would also have to be taken into account. Alternatively, a combination of single- and mixed-cell cultures and mathematical modelling may be able to identify the nature of interactions *in vitro* and provide some useful insights into the co-evolution of EGFR and PDGFRA sub-populations *in vivo* if such data were to become available.

## Acknowledgments

Authors KRS, NLT, LSH, LC, and AHD gratefully acknowledge funding for this work through NCI U01CA220378.

BM, MEH, RR, SJS, DA and MRO gratefully acknowledge PhD studentship funding from the UK EPSRC (reference EP/N50970X/1).

## References

[1] R. Stupp, W. P. Mason, M. J. Van Den Bent, M. Weller, B. Fisher, M. J. Taphoorn, et al. “Radiotherapy plus concomitant and adjuvant temozolomide for glioblastoma”. In: New England Journal of Medicine 352.10 (2005), pp. 987–996.

[2] R. Stupp, M. E. Hegi, W. P. Mason, M. J. Van Den Bent, M. J. Taphoorn, R. C. Janzer, et al. “Effects of radiotherapy with concomitant and adjuvant temozolomide versus radiotherapy alone on survival in glioblastoma in a randomised phase III study: 5-year analysis of the EORTC-NCIC trial”. In: The lancet oncology 10.5 (2009), pp. 459–466.

[3] R. Bonavia, W. K. Cavenee, and F. B. Furnari. “Heterogeneity maintenance in glioblastoma: a social network”. In: Cancer research 71.12 (2011), pp. 4055–4060.

[4] Z. An, O. Aksoy, T. Zheng, Q.-W. Fan, and W. A. Weiss. “Epidermal growth factor receptor and EGFRvIII in glioblastoma: signaling pathways and targeted therapies”. In: Oncogene 37.12 (2018), pp. 1561–1575.

[5] M. Nakada, D. Kita, T. Watanabe, Y. Hayashi, and J.-i. Hamada. “The mechanism of chemoresistance against tyrosine kinase inhibitors in malignant glioma”. In: Brain tumor pathology 31.3 (2014), pp. 198–207.

[6] N. J. Szerlip, A. Pedraza, D. Chakravarty, M. Azim, J. McGuire, Y. Fang, et al. “Intratumoral heterogeneity of receptor tyrosine kinases EGFR and PDGFRA amplification in glioblastoma defines subpopulations with distinct growth factor response”. In: Proceedings of the National Academy of Sciences 109.8 (2012), pp. 3041–3046.

[7] L. S. Hu, S. Ning, J. M. Eschbacher, L. C. Baxter, N. Gaw, S. Ranjbar, et al. “Radiogenomics to characterize regional genetic heterogeneity in glioblastoma”. In: Neuro-oncology 19.1 (2017), pp. 128–137.

[8] A. Sottoriva, I. Spiteri, S. G. Piccirillo, A. Touloumis, V. P. Collins, J. C. Marioni, et al. “Intratumor heterogeneity in human glioblastoma reflects cancer evolutionary dynamics”. In: Proceedings of the National Academy of Sciences 110.10 (2013), pp. 4009–4014.

[9] S. J. Smith, M. Diksin, S. Chhaya, S. Sairam, M. A. Estevez-Cebrero, and R. Rahman. “The invasive region of glioblastoma defined by 5ALA guided surgery has an altered cancer stem cell marker profile compared to central tumour”. In: International journal of molecular sciences 18.11 (2017), p. 2452.

[10] J. G. Lyons, E. Lobo, A. M. Martorana, and M. R. Myerscough. “Clonal diversity in carcinomas: its implications for tumour progression and the contribution made to it by epithelial-mesenchymal transitions”. In: Clinical & experimental metastasis 25.6 (2008), pp. 665–677.

[11] C. Lopez-Gines, R. Gil-Benso, R. Ferrer-Luna, R. Benito, E. Serna, J. Gonzalez-Darder, et al. “New pattern of EGFR amplification in glioblastoma and the relationship of gene copy number with gene expression profile”. In: Modern Pathology 23.6 (2010), pp. 856–865.

[12] F. B. Furnari, T. F. Cloughesy, W. K. Cavenee, and P. S. Mischel. “Heterogeneity of epidermal growth factor receptor signalling networks in glioblastoma”. In: Nature Reviews Cancer 15.5 (2015), p. 302.

[13] B. R. Voldborg, L. Damstrup, M. Spang-Thomsen, and H. S. Poulsen. “Epidermal growth factor receptor (EGFR) and EGFR mutations, function and possible role in clinical trials”. In: Annals of Oncology 8.12 (1997), pp. 1197–1206.

[14] J. J. Parker, K. R. Dionne, R. Massarwa, M. Klaassen, N. K. Foreman, L. Niswander, et al. “Gefitinib selectively inhibits tumor cell migration in EGFR-amplified human glioblastoma”. In: Neuro-oncology 15.8 (2013), pp. 1048–1057.

[15] K. M. Talasila, A. Soentgerath, P. Euskirchen, G. V. Rosland, J. Wang, P. C. Huszthy, et al. “EGFR wild-type amplification and activation promote invasion and development of glioblastoma independent of angiogenesis”. In: Acta neuropathologica 125.5 (2013), pp. 683–698.

[16] N. Shinojima, K. Tada, S. Shiraishi, T. Kamiryo, M. Kochi, H. Nakamura, et al. “Prognostic value of epidermal growth factor receptor in patients with glioblastoma multiforme”. In: Cancer research 63.20 (2003), pp. 6962–6970.

[17] A. Alentorn, Y. Marie, C. Carpentier, B. Boisselier, M. Giry, M. Labussiere, et al. “Prevalence, clinicopathological value, and co-occurrence of PDGFRA abnormalities in diffuse gliomas”. In: Neurooncology 14.11 (2012), pp. 1393–1403.

[18] P. Blume-Jensen and T. Hunter. “Oncogenic kinase signalling”. In: Nature 411.6835 (2001), p. 355.

[19] C. G. A. R. Network et al. “Comprehensive genomic characterization defines human glioblastoma genes and core pathways”. In: Nature 455.7216 (2008), p. 1061.

[20] M. Lino and A. Merlo. “PI3Kinase signaling in glioblastoma”. In: Journal of neuro-oncology 103.3 (2011), pp. 417–427.

[21] M. Snuderl, L. Fazlollahi, L. P. Le, M. Nitta, B. H. Zhelyazkova, C. J. Davidson, et al. “Mosaic amplification of multiple receptor tyrosine kinase genes in glioblastoma”. In: Cancer cell 20.6 (2011), pp. 810–817.

[22] F. Chen and L. Ding. “Co-survival of the fittest few: mosaic amplification of receptor tyrosine kinases in glioblastoma”. In: Genome biology 13.1 (2012), p. 141.

[23] M. J. Borad, M. D. Champion, J. B. Egan, W. S. Liang, R. Fonseca, A. H. Bryce, et al. “Integrated genomic characterization reveals novel, therapeutically relevant drug targets in FGFR and EGFR pathways in sporadic intrahepatic cholangiocarcinoma”. In: PLoS genetics 10.2 (2014).

[24] D. W. Craig, J. A. O’Shaughnessy, J. A. Kiefer, J. Aldrich, S. Sinari, T. M. Moses, et al. “Genome and transcriptome sequencing in prospective metastatic triple-negative breast cancer uncovers therapeutic vulnerabilities”. In: Molecular cancer therapeutics 12.1 (2013), pp. 104–116.

[25] R. Mehrian-Shai, M. Yalon, I. Moshe, I. Barshack, D. Nass, J. Jacob, et al. “Identification of genomic aberrations in hemangioblastoma by droplet digital PCR and SNP microarray highlights novel candidate genes and pathways for pathogenesis”. In: BMC genomics 17.1 (2016), p. 56.

[26] J. Alfonso, K. Talkenberger, M. Seifert, B. Klink, A. Hawkins-Daarud, K. Swanson, et al. “The biology and mathematical modelling of glioma invasion: a review”. In: Journal of the Royal Society Interface 14.136 (2017), p. 20170490.

[27] K. Swanson, R. Rostomily, and E. Alvord Jr. “A mathematical modelling tool for predicting survival of individual patients following resection of glioblastoma: a proof of principle”. In: British journal of cancer 98.1 (2008), pp. 113–119.

[28] K. Swanson, R. Rostomily, and E. Alvord Jr. “Confirmation of a theoretical model describing the relative contributions of net growth and dispersal in individual infiltrating gliomas”. In: Can J Neurol Sci 30 (2003), pp. 407–434.

[29] K. R. Swanson, E. C. Alvord Jr, and J. Murray. “A quantitative model for differential motility of gliomas in grey and white matter”. In: Cell proliferation 33.5 (2000), pp. 317–329.

[30] K. R. Swanson, E. C. Alvord, and J. Murray. “Virtual brain tumours (gliomas) enhance the reality of medical imaging and highlight inadequacies of current therapy”. In: British journal of cancer 86.1 (2002), pp. 14–18.

[31] K. R. Swanson, C. Bridge, J. Murray, and E. C. Alvord Jr. “Virtual and real brain tumors: using mathematical modeling to quantify glioma growth and invasion”. In: Journal of the neurological sciences 216.1 (2003), pp. 1–10.

[32] K. Swanson, H. Harpold, D. Peacock, R. Rockne, C. Pennington, L. Kilbride, et al. “Velocity of radial expansion of contrast-enhancing gliomas and the effectiveness of radiotherapy in individual patients: a proof of principle”. In: Clinical oncology 20.4 (2008), pp. 301–308.

[33] A. L. Baldock, S. Ahn, R. Rockne, S. Johnston, M. Neal, D. Corwin, et al. “Patient-specific metrics of invasiveness reveal significant prognostic benefit of resection in a predictable subset of gliomas”. In: PLoS One 9.10 (2014).

[34] P. R. Jackson, J. Juliano, A. Hawkins-Daarud, R. C. Rockne, and K. R. Swanson. “Patient-specific mathematical neuro-oncology: using a simple proliferation and invasion tumor model to inform clinical practice”. In: Bulletin of mathematical biology 77.5 (2015), pp. 846–856.

[35] C. H. Wang, J. K. Rockhill, M. Mrugala, D. L. Peacock, A. Lai, K. Jusenius, et al. “Prognostic significance of growth kinetics in newly diagnosed glioblastomas revealed by combining serial imaging with a novel biomathematical model”. In: Cancer research 69.23 (2009), pp. 9133–9140.

[36] K. J. Painter and T. Hillen. “Volume-filling and quorum-sensing in models for chemosensitive movement”. In: Can. Appl. Math. Quart 10.4 (2002), pp. 501–543.

[37] P. Gerlee and S. Nelander. “The impact of phenotypic switching on glioblastoma growth and invasion”. In: PLoS computational biology 8.6 (2012).

[38] K. R. Swanson, R. C. Rockne, J. Claridge, M. A. Chaplain, E. C. Alvord, and A. R. Anderson. “Quantifying the role of angiogenesis in malignant progression of gliomas: in silico modeling integrates imaging and histology”. In: Cancer research 71.24 (2011), pp. 7366–7375.

[39] A. Hawkins-Daarud, S. K. Johnston, and K. R. Swanson. “Quantifying uncertainty and robustness in a biomathematical model–based patient-specific response metric for glioblastoma”. In: JCO clinical cancer informatics 3 (2019), pp. 1–8.

[40] C. A. Smith and C. A. Yates. “The auxiliary region method: a hybrid method for coupling PDE-and Brownian-based dynamics for reaction–diffusion systems”. In: Royal Society open science 5.8 (2018), p. 180920.

