## Supplementary Material for "Identifying the spatial and temporal dynamics of molecularly-distinct glioblastoma sub-populations"

Here we present additional figures exploring the effects of various selection advantages and the timing and positioning of EGFR and PDGFRA amplified sub-population introductions on the amplification patterns we see in our simulated tumours. These figures show the general trends in changes to the proportions of simulations with neither gene, only the EGFR gene, only the PDGFRA gene and both genes amplified that changing each of these factors produces. We note that the effects of changing each of these factors are symmetric with respect to the proportions of simulations with only EGFR and only PDGFRA amplified, e.g. giving $E$ cells a 50% proliferative advantage and $P$ cells no advantages, produces the same simulation proportions of neither and both amplified cells as giving the $P$ population this advantage and $E$ no advantage with the proportion with only EGFR amplified reflected down the only PDGFRA amplified axis.

All of the following figures are produced from simulations with the same parameters and assumptions outlined in Section 3.3, apart from where parameter differences are indicated in the figure captions.

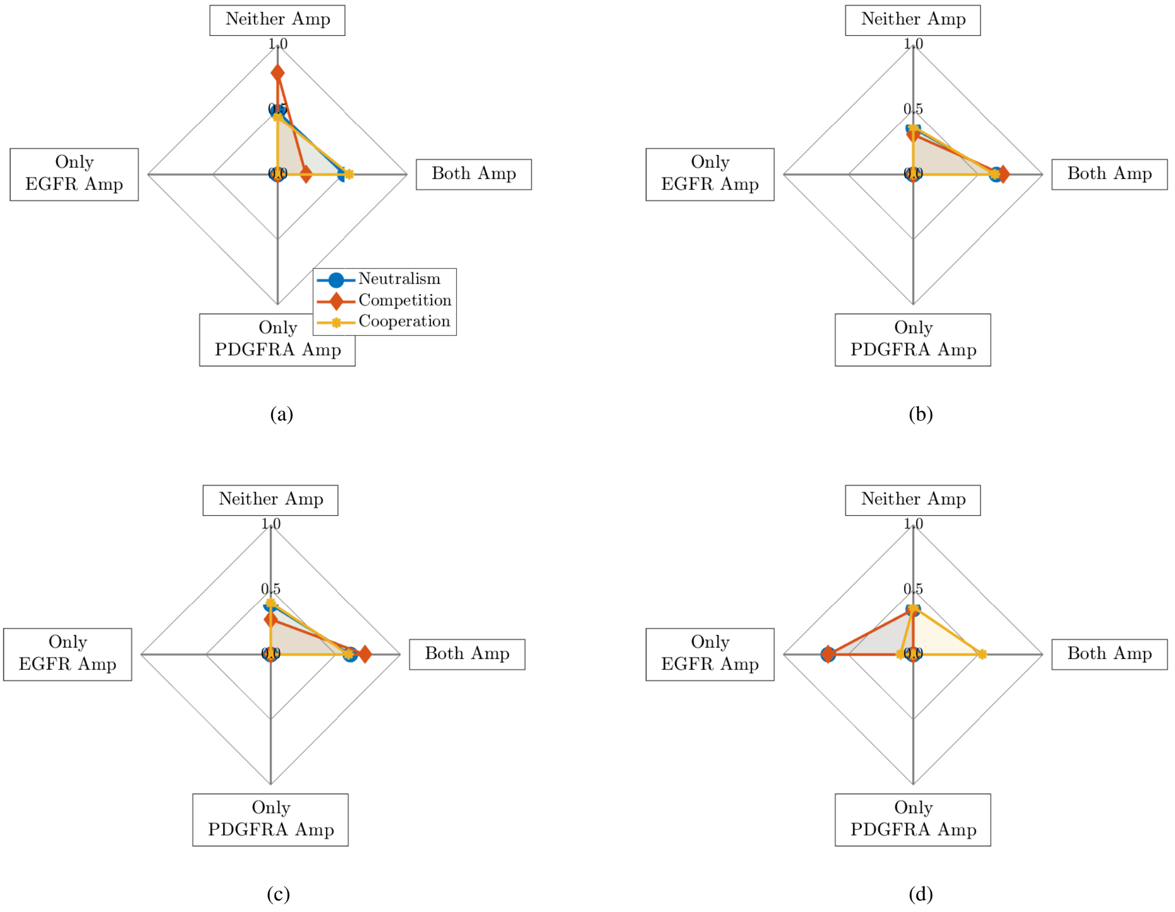


Figure 1: Plot showing the mean proportions of simulations with neither gene (Neither Amp), only the EGFR gene (Only EGFR Amp), only the PDGFRA gene (Only PDGFRA Amp) and both genes (Both Amp) amplified under different interactions when the $E$ and $P$ sub-populations are given various selection advantages: (a) EGFR 50% invasive advantage, PDGFRA 50% invasive advantage ($\rho_E = \rho_P = \rho_N$ and $D_E = D_P = 1.5D_N$); (b) EGFR 50% proliferative advantage, PDGFRA 50% proliferative advantage ($\rho_E = \rho_P = 1.5\rho_N$ and $D_E = D_P = D_N$); (c) EGFR 50% proliferative and invasive advantage, PDGFRA 50% proliferative and invasive advantage ($\rho_E = \rho_P = 1.5\rho_N$ and $D_E = D_P = 1.5D_N$); (d) EGFR 50% proliferative advantage, PDGFRA 50% invasive advantage ($\rho_E = 1.5\rho_P = 1.5\rho_N$ and $1.5D_E = D_P = 1.5D_N$).

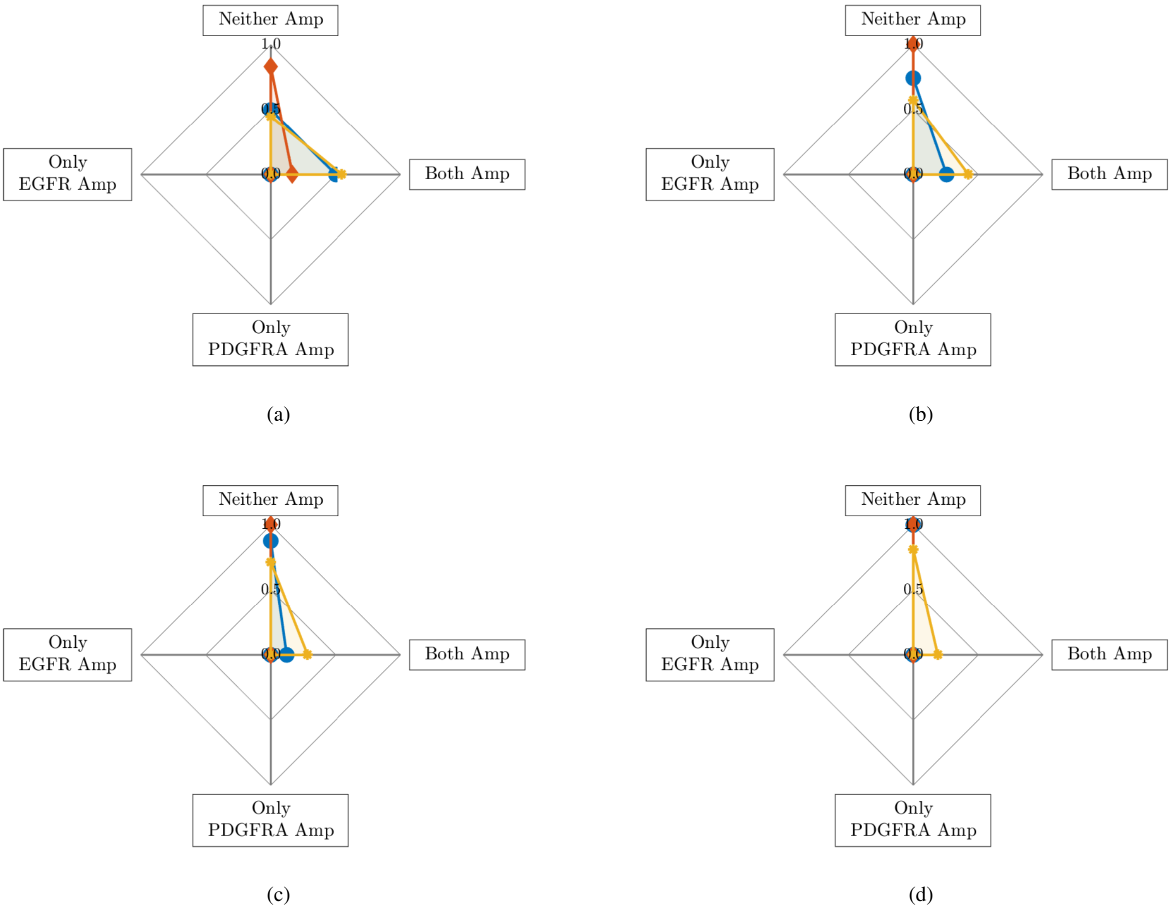


Figure 2: Plot showing the mean proportions of simulations with neither gene (Neither Amp), only the EGFR gene (Only EGFR Amp), only the PDGFRA gene (Only PDGFRA Amp) and both genes (Both Amp) amplified under different interactions when the $E$ and $P$ sub-populations are introduced at the same time which changes: (a) $t_E^* = t_P^* = t_1^*$; (b) $t_E^* = t_P^* = t_3^*$; (c) $t_E^* = t_P^* = t_5^*$; (d) $t_E^* = t_P^* = t_7^*$.

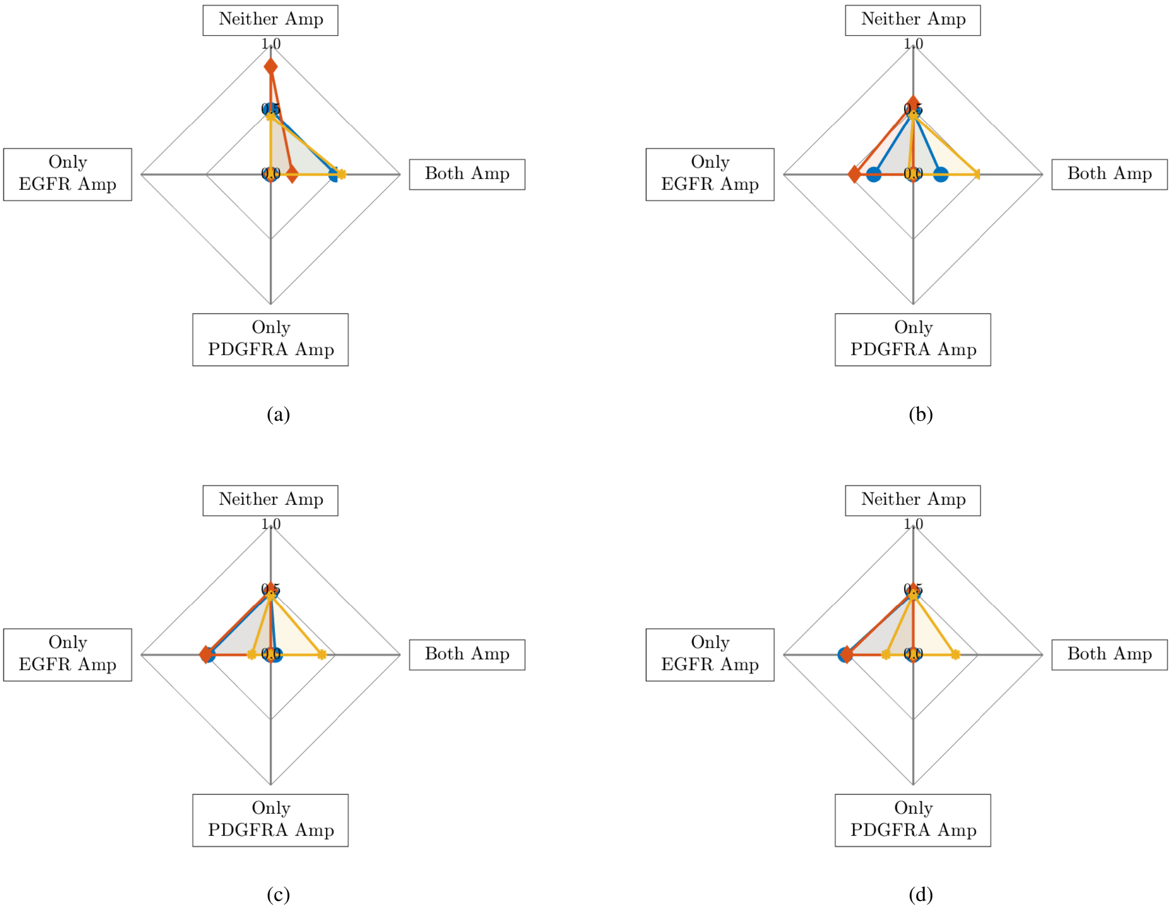


Figure 3: Plot showing the mean proportions of simulations with neither gene (Neither Amp), only the EGFR gene (Only EGFR Amp), only the PDGFRA gene (Only PDGFRA Amp) and both genes (Both Amp) amplified under different interactions when the $E$ population is introduced at $t_E^* = t_1^*$ and $P$ is introduced at: (a) $t_P^* = t_1^*$; (b) $t_P^* = t_3^*$; (c) $t_P^* = t_5^*$; (d) $t_P^* = t_7^*$.

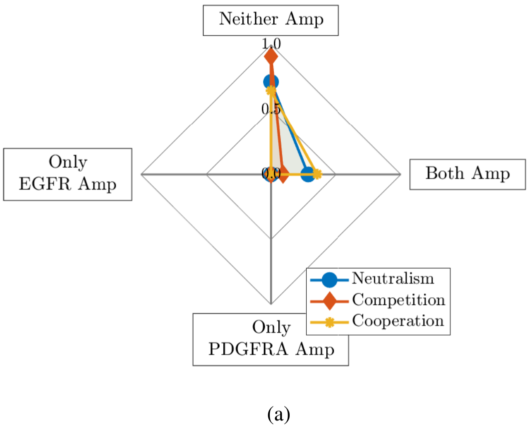


Figure 4: Plot showing the mean proportions of simulations with neither gene (Neither Amp), only the EGFR gene (Only EGFR Amp), only the PDGFRA gene (Only PDGFRA Amp) and both genes (Both Amp) amplified under different interactions when the $E$ and $P$ populations are introduced at the same location, which changes: (a) $x_E^* = x_P^* = x_1^*$; (b) $x_E^* = x_P^* = x_2^*$; (c) $x_E^* = x_P^* = x_3^*$; (d) $x_E^* = x_P^* = x_4^*$.

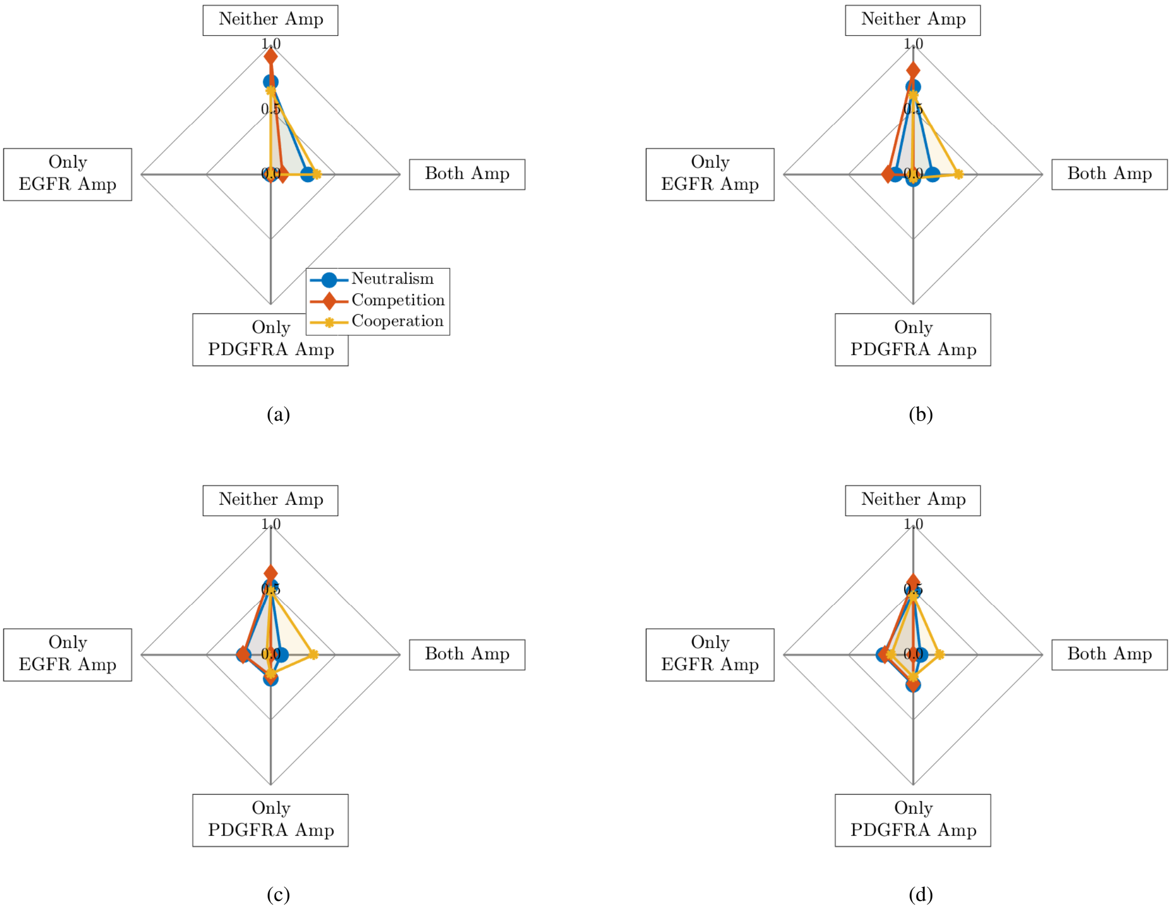


Figure 5: Plot showing the mean proportions of simulations with neither gene (Neither Amp), only the EGFR gene (Only EGFR Amp), only the PDGFRA gene (Only PDGFRA Amp) and both genes (Both Amp) amplified under different interactions when the $E$ population is introduced at $x_E^* = x_1^*$ and $P$ is introduced at: (a) $x_P^* = x_1^*$; (b) $x_P^* = x_3^*$; (c) $x_P^* = x_5^*$; (d) $x_P^* = x_7^*$.
